# Pathogen effector-associated nuclear-localization of RPW8.2 amplifies its expression to boost immunity in Arabidopsis

**DOI:** 10.1101/2022.07.28.501905

**Authors:** Jing-hao Zhao, Yan-yan Huang, He Wang, Xue-mei Yang, Jing Fan, Yan Li, Mei Pu, Shi-xin Zhou, Ji-wei Zhang, Zhi-xue Zhao, Guo-bang Li, Beenish Hassan, Xiao-hong Hu, Xue-wei Chen, Shunyuan Xiao, Xian-jun Wu, Wen-ming Wang

**Author notes:** Authors for correspondence: Wen-Ming Wang Xian-Jun Wu Shunyuan Xiao. **One-sentence summary:** The powdery mildew pathogen *G. cichoracearum* secretes GcR8IP1 protein to increase nuclear localization of RPW8.2 and promote transcriptional amplification of *RPW8.2* to boost immunity in Arabidopsis.

## Abstract

RESISTANCE TO POWDERY MILDEW 8 (RPW8) defines a unique N-terminal coiled-coil domain of nucleotide-binding and leucine-rich repeat immune receptors required for immune signaling in plants. Arabidopsis RPW8.2 is specifically induced by the powdery mildew (PM) fungus (*Golovinomyces cichoracearum*) in the infected epidermal cells to activate immunity. The mechanism of RPW8.2-induction is not well understood. Here, we identify a *G. cichoracearum* factor delivered to the nucleus of the host cell, named Gc-RPW8.2 interacting protein 1 (GcR8IP1). Ectopic expression of GcR8IP1 in Arabidopsis or *Nicotiana benthamiana* suppressed host immune responses and enhanced susceptibility to PM. Host-induced gene silencing of *GcR8IP1* compromised PM infectivity in susceptible Arabidopsis plants. Co-expression of GcR8IP1 with RPW8.2 increased nuclear localization of RPW8.2, which in turn, promoted transcriptional amplification of *RPW8.2*. Thus, RPW8.2-dependent defense strengthening is due to altered partitioning of RPW8.2 by an effector of a PM fungus, which exemplifies an atypical form of effector-triggered immunity.

## Introduction

Plants mount a two-tiered immune system that consists of pattern-triggered immunity (PTI) and effector-triggered immunity (ETI) against microbial pathogens (Wang et al., 2009; Jones and Dangl, 2006). PTI and ETI are mutually potentiated upon their activations by the recognitions of the cell-surface and intracellular immune receptors with the conserved pathogen-associated molecular patterns (PAMPs) and pathogen effectors, respectively (Ngou et al., 2021; Yuan et al., 2021). In many cases, intracellular immune receptors belong to the nucleotide-binding leucine-rich repeat (NLR) proteins that are classified into two broad classes based on their N-terminal domains: Toll and interleukin-1 receptor (TIR) NLRs (TNLs) and coiled-coil (CC) NLRs (CNLs) (Kourelis and Van Der Hoorn, 2018). The TIR domains of TNLs have been demonstrated to possess NADase activity that is mutually exclusive with their 2’,3’-cAMP/cGMP synthetase activity, both of which are required for the activation of immune response (Horsefield et al., 2019; Wan et al., 2019; Yu et al., 2022). The CNL ZAR1 resistosome acts as a Ca^2+^-permeable cation channel in the plasma membrane relaying immune signaling (Wang et al., 2019; Bi et al., 2021). Within CNL proteins, one basal clade is distinguished by having a CC domain resembling the *Arabidopsis thaliana* RESISTANCE TO POWDERY MILDEW 8 (RPW8) proteins, denoted as the CC_R_ domain (Collier et al., 2011). The CC_R_ CNLs include two related helper NLRs, namely N-Required Gene 1 (NRG1) and Activated Disease Resistance 1 (ADR1) (Roberts et al., 2013; Peart et al., 2005). In contrast to the CC domain of canonical CNL proteins, the CC_R_ domains of both NRG1 and ADR1 family proteins are sufficient for the induction of defense responses (Collier et al., 2011). Besides, the ADR1 and NRG1 families play an unequally redundant role in immune signaling transduction of TNL- and CNL-mediated ETI (Dong et al., 2016; Castel et al., 2019; Wu et al., 2019; Saile et al., 2020). These two unique families of NLR proteins have also been termed as RNLs because of their N-terminal CC_R_ domain homologous to the RPW8 family proteins, which include RPW8.1, RPW8.2, and a few homologues of RPW8 (Dong et al., 2016; Castel et al., 2019; Collier et al., 2011; Xiao et al., 2004). Both RPW8.1 and RPW8.2 confer broad-spectrum resistance to powdery mildew (PM) (Xiao et al., 2003; Wang et al., 2009; Ma et al., 2014; Xiao et al., 2001).

PM fungi are obligate biotrophic pathogens that require a living host to complete their lifecycle (Lipka et al., 2008). PM infection induces reorganizations in host cell structure and changes in cell physiology (Hückelhoven and Panstruga, 2011). Particularly, PM fungi establish haustoria in epidermal cells of their hosts to steal water and nutrients for their proliferation (Lipka et al., 2008). The haustoria of PM are encased by the extra-haustorial membrane (EHM) that forms the host-pathogen interface and the battle frontiers (Kwaaitaal et al., 2017). RPW8.2 is targeted to the EHM via the VAMP721/722-mediated vesicle trafficking pathway upon the invasion of PM fungi that highly induce the expression of *RPW8.2* (Wang et al., 2009; Kim et al., 2014). RPW8.2 is also partitioned into the nucleus and the cytoplasm to mount effective defense responses, such as the deposition of callose to encase the haustorial complex and the enrichment of H_2_O_2_ in the haustorial complex (Wang et al., 2009; Huang et al., 2019). Both RPW8.1 and RPW8.2 engage EDS1-dependent and SA signaling to trigger a positive transcriptional amplification circuit (Xiao et al., 2003, 2005). Besides, *RPW8.2* is strongly and specifically induced in epidermal cells invaded by the adapted PM isolate *Golovinomyces cichoracearum* (*Gc*) UCSC1 (Xiao et al., 2003). However, it remains unclear how the highly cell-type-specific induction of *RPW8.2* is achieved.

To explore how RPW8.2 activates defense and how RPW8.2 is regulated, we performed a yeast-two-hybridization screen and identified a candidate protein from *Gc* USCS1, designated Gc-RPW8.2 interacting protein 1 (GcR8IP1). We demonstrated that GcR8IP1 is a secreted effector that plays a crucial role in promoting *Gc* UCSC1’s virulence. GcR8IP1 is nucleus-localized. Its binding to RPW8.2 increases RPW8.2’s nuclear localization where RPW8.2 activates a transcriptional self-amplification in haustorium-invaded epidermal cells. Thus, our data provide insight into the mechanism of RPW8.2-mediated immunity against PM as an atypical form of effector-triggered immunity.

## Results

### RPW8.2 interacts with a powdery mildew protein GcR8IP1

In a yeast two-hybridization screen, we identified a candidate RPW8.2-interacting protein (R8IP) predicted to be 129 amino acids (aa) of a protein encoded by a gene in the *Gc* UCSC1 genome (Figure 1A). The full length message RNA (mRNA) was consistent with the reported sequences (GenBank: RKF55523.1) (Wu et al., 2018), here designated *GcR8IP1*. *GcR8IP1* encodes a 352 aa protein that is predicted as a cytoplasmic fungal effector by EffectorP (Sperschneider and Dodds, 2022), containing a presumable N-terminal secretion signal peptide (SP, which was not predicted using SignaIP 6.0 (Teufel et al., 2022)), followed by a REALLY INTERESTING NEW GENE (RING) domain and a MENAGE A TROIS 1 (MAT1) domain, and a C-terminus with unknown feature (Figure 1B). GcR8IP1 is highly conserved in all PM proteomes predicted from currently available PM genome sequences (Supplemental Figure S1). The 129 aa (hereon GcR8IP1-M1) identified in the Y2H screen contains the SP-RING and partial MAT1.

**Figure 1.**
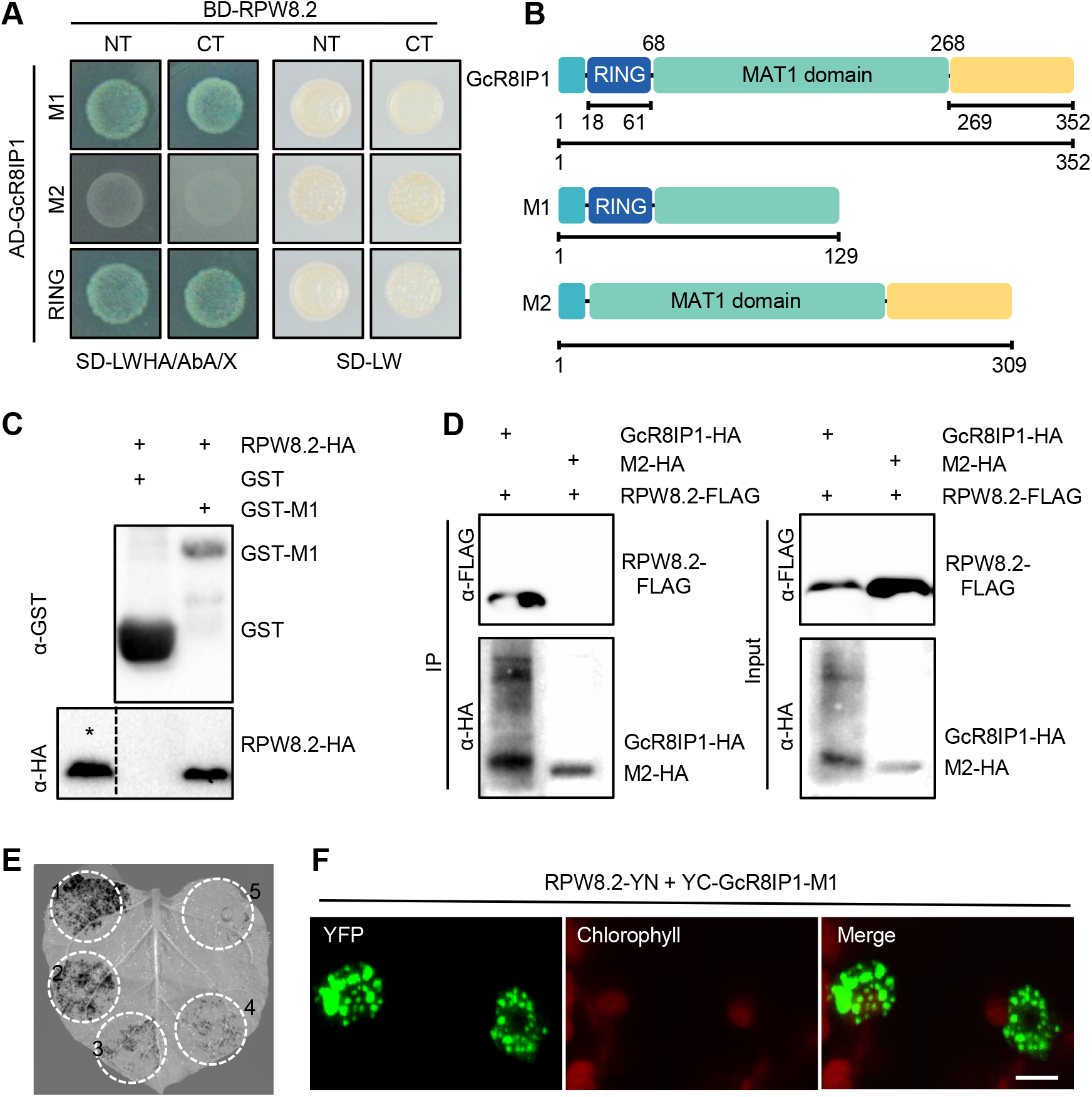
RPW8.2 interacts with GcR8IP1. A, Yeast-two-hybrid (Y2H) assays show the associations of the first 100 amino acid residues (aa) (NT) and the 101-174 aa (CT) of RPW8.2 with the first 129 aa (M1) and the RING domain (RING) of GcR8IP1. The NT and CT of RPW8.2 fused with the DNA-binding domain (BD) of GAL4 were used as baits and the indicated portion of GcR8IP1 fused with the transcriptional activation domain (AD) of GAL4 were used as preys. GcR8IP1-M2 is a RING domain-deleted mutant. Transformed yeast was grown on selective medium (SD) lacking adenine, histidine, leucine, and tryptophan with AbA and X-α-Gal (SD-LWHA/AbA/X). SD-LW indicates SD lacks Leu and Trp. B, A schematic diagram shows the position of each domain and deletions in GcR8IP1. The position of the RING finger domain, MAT1 domain, and unknown motif (269-352) are indicated. Numbers indicate amino acid positions. C, A semi-*in vivo* GST pull-down assay shows the interaction of RPW8.2-HA with GST-GcR8IP1-M1 (GST-M1). GST or GST-M1 immobilized on GST beads was co-incubated with lysates of *Nicotiana benthamiana* leaves transiently expressing 35S_pro_:RPW8.2-HA. The beads were washed and pelleted for immunoblotting with an anti-GST antibody and an anti-HA antibody. The input lysates were added as expression control of 35S_pro_:RPW8.2-HA (*) in the immunoblotting with an anti-HA antibody. D, Co-immunoprecipitation assay shows the interaction of RPW8.2-FLAG with GcR8IP1-HA. Total proteins were extracted from *N. benthamiana* leaves co-expressing 35S_pro_:RPW8.2-FLAG with 35S_pro_:GcR8IP1-HA or 35S_pro_:GcR8IP1-M2-HA (M2-HA). Protein complexes were pulled down using an anti-HA antibody conjugated on agarose beads and the co-precipitated complex was examined by western blotting using an anti-FLAG antibody and an anti-HA antibody. E, Luciferase complementation imaging (LCI) assay. Transiently expression was conducted via co-infiltration of the Agrobacterial strains carrying constructs for expressing NLuc-Rac1 and CLuc-OsRbohB (1) as positive control, NLuc-RPW8.2-NT and CLuc-GcR8IP1 (2), NLuc-RPW8.2-CT and CLuc-GcR8IP1 (3), NLuc-RPW8.2 and CLuc-GcR8IP1 (4), and NLuc-RPW8.2-NT and CLuc-GcR8IP1-M2 (5). LCI images were acquired from *N. benthamiana* leaves at 48 hours post infiltration. F, Bimolecular fluorescence complementation assay shows the association of RPW8.2 and GcR8IP1-M1 in the nucleus. 35S_pro_:RPW8.2-YN and 35S_pro_:YC-GcR8IP1-M1 were transiently co-expressed in leaves of 6-week-old *N. benthamiana* plants and examined by confocal microscopy. Size bar, 10 μm.

RPW8.2 has two nuclear localization signals (NLSs) in its first 100 aa (RPW8.2-NT) and two nuclear export signals in the 101-174 aa (RPW8.2-CT) (Huang et al., 2014, 2019). Both RPW8.2-NT and RPW8.2-CT interacted with GcR8IP1-M1 and the RING domain (18-61 aa), but not with the RING-deleted mutant (M2), of GcR8IP1 in yeast (Figure 1A). RPW8.2-HA was pulled down by GST-GcR8IP1-M1 but not by GST (Figure 1B, C). The association of RPW8.2 with full length GcR8IP1 was confirmed by co-immunoprecipitation (Co-IP) and LCI (Firefly Luciferase Complementation Imaging) assays in *N. benthamiana* (Figure 1D-F). BiFC assays showed that RPW8.2-YN and YC-GcR8IP1-M1 formed puncta in the nucleus and the RING domain of GcR8IP1 is sufficient for the interaction with RPW8.2 (Figure 1F and Supplemental Figure S2). These data indicate that RPW8.2 interacts with GcR8IP1 in the nucleus.

### GcR8IP1 contains a secretion signal peptide and is delivered into the host nucleus

We tested if the presumable SP of GcR8IP1 could restore secretion of invertase in yeast using the invertase secretion-deficient yeast strain YTK12 from a previous report (Jacobs et al., 1997). The first 16, 26 and 40 aa of GcR8IP1 restored the invertase secretion-deficient mutant yeast strain YTK12 to grow on the YPRAA medium as the positive control SP of Avr1b from a previous report (Fang et al., 2016), indicating properly secretion of invertase (Figure 2A). TTC (2,3,5-triphenyl-2-tetrazolium chloride) assay confirmed that the SP of Avr1b and the first 16, 26, and 40 aa of GcR8IP1 enabled the secretion of invertase that catalyzes TTC into red insoluble substance, but not the negative control of Mg87 (Figure 2A). Consistently, the first 40 aa of GcR8IP1 brought GFP into the apoplast space as a fusion protein when transiently expressed in leaf epidermal cells of *N. benthamiana*, indicating secretion *in planta* (Figure 2B). These results suggest that GcR8IP1 may be a SP-containing effector protein from *Gc* UCSC1.

**Figure 2.**
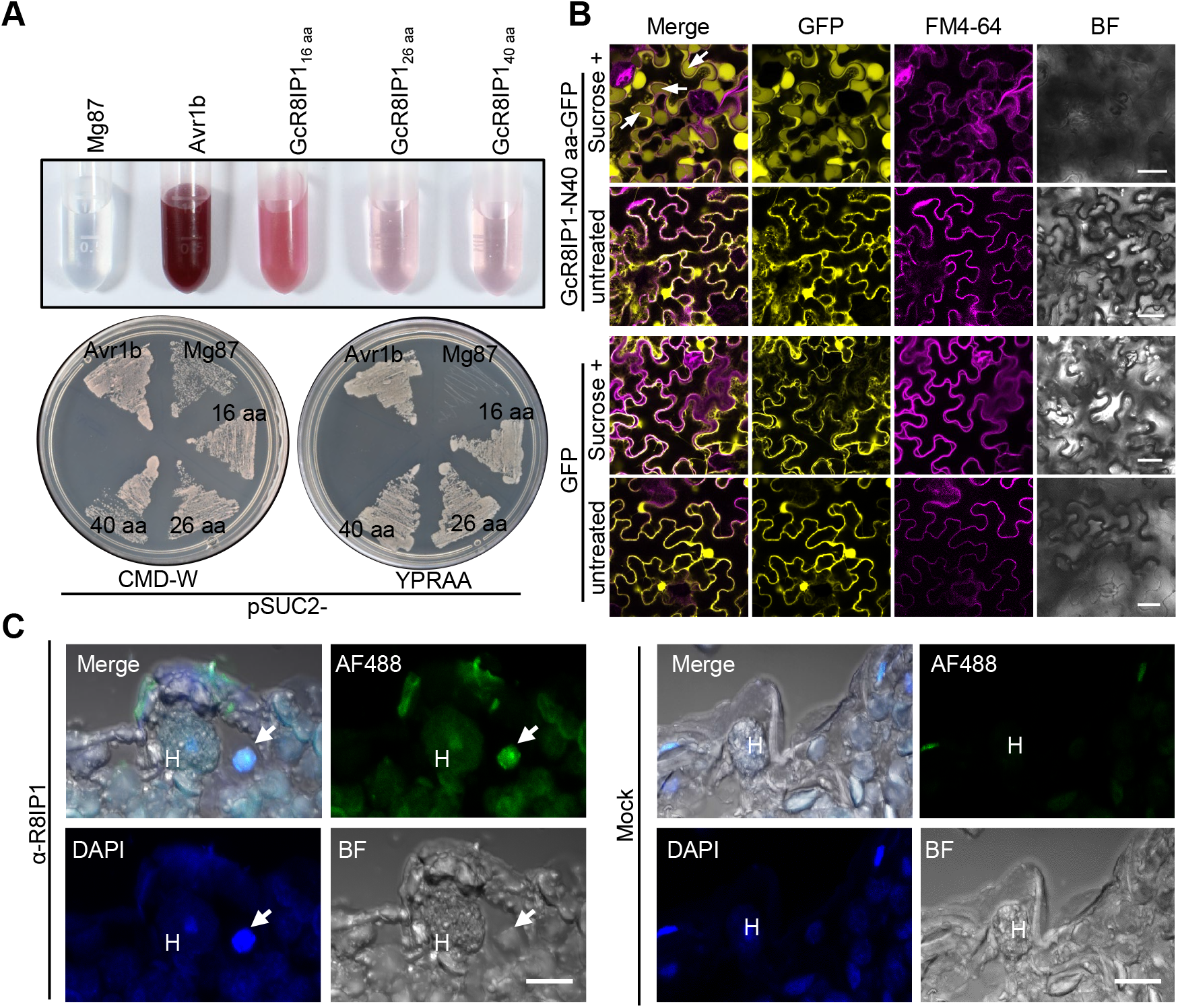
GcR8IP1 is a secretory protein. A, Validation of the GcR8IP1 signal peptide by yeast invertase secretion assay. The DNA fragment encoding the first 16 aa, 26 aa or 40 aa of GcR8IP1 was in-frame fused to yeast mature invertase sequence in the pSUC2 vector and expressed in YTK12. A functional signal peptide enables yeast growth on both CMD-W and YPRAA, and also reduces TTC (2,3,5-triphenyl tetrazolium chloride) into a red formazan. The N-terminal sequence of Mg87 and the signal peptide of Avr1b were used as negative and positive control, respectively. B, Confocal images show the validation of the GcR8IP1 signal peptide. *N. benthamiana* leaves transiently expressing the first 40 aa of GcR8IP1 fused with GFP (GcR8IP1-N40 aa-GFP) or GFP, respectively, were stained with FM4-64 and imaged at 48 hpi. Note that GcR8IP1-N40 aa-GFP was secreted into the apoplast (arrows). Sucrose +: samples were incubated with 12% sucrose for 10 min for plasmolysis. Size bars, 25 μm. C, Immunofluorescence staining images show the localization of GcR8IP1 in the nucleus (arrow). The slides were prepared from Arabidopsis *pad4-1 sid2-1* leaves 3 days post inoculation of *G. cichoracearum* UCSC1. The subcellular localization of GcR8IP1 (green) was detected by a primary antibody raised against GcR8IP1 and visualized by Alexa Fluor^®^ 488-conjugated secondary antibodies (AF488) that binds the primary antibody. The nucleus (blue) was counterstained with 4’,6-diamidino-2-phenylindole (DAPI). H: haustoria. α-R8IP1: antibody of GcR8IP1. Mock: rabbit antisera. Size bar, 10 μm.

The expression of *GcR8IP1* was up-regulated at 3 days post inoculation (dpi), 4 dpi, and 6 dpi of *Gc* UCSC1 on rosette leaves of *Arabidopsis* (Supplemental Figure S3A). The fungal biomass maintained at a low amount and showed no significant difference within 6 dpi, but had a sharp increase at 7 dpi and 12-13 dpi (Supplemental Figure S3B). It is well-known that the establishment of haustoria starts at 24 hours post inoculation and reached a summit at 3-4 dpi, events that support the proliferation and sporulation of PM at 5-6 dpi (Koh et al., 2005; Micali et al., 2008) . Therefore, the expression pattern of *GcR8IP1* may be associated with the establishment of the haustoria and sporulation of PM.

Next, we tried to express GcR8IP1-GFP in transgenic *Magnaporthe oryzae* to test its secretion into the host cell during infection following a previous report (Zhang et al., 2020). Unfortunately, we failed to detect GFP in the positive transgenic *M*. *oryzae* strains during infection of rice. Then, we made a polyclonal antibody for GcR8IP1 to examine its subcellular localization during infection via immunofluorescence assay. While the mock treatment generated a background signal, more intense signals were observed in the nuclei of both the haustorium and the host cell as puncta than in the other organelles (Figure 2C), indicating that GcR8IP1 is delivered into the host cell and localized in the nucleus. Consistently, GcR8IP1-GFP was localized in the nucleus when it was transiently expressed in *N*. *benthamiana* (Supplemental Figure S4). These data indicate that GcR8IP1 is secreted into host cells and localized in the nucleus.

### Ectopic expression of GcR8IP1 suppresses immune responses

Transient expression of the RPW8.2 C-terminal (CC_RPW8.2,_ RPW8.2 101-174 aa) triggers cell death in *N. benthamiana* (Huang et al., 2019). Such cell death was observed at 5 days, but not at 60 hours post infiltration in *N. benthamiana* when CC_RPW8.2_ was transiently co-expressed with GcR8IP1 via agroinfiltration assay, indicating CC_RPW8.2_-triggered cell death was delayed (Figure 3A). RPW8.2-HA-triggered ion leakage was significantly reduced when it was transiently co-expressed with GcR8IP1 (Figure 3B, C). These data indicate GcR8IP1 may suppress RPW8.2-meidated immunity.

**Figure 3.**
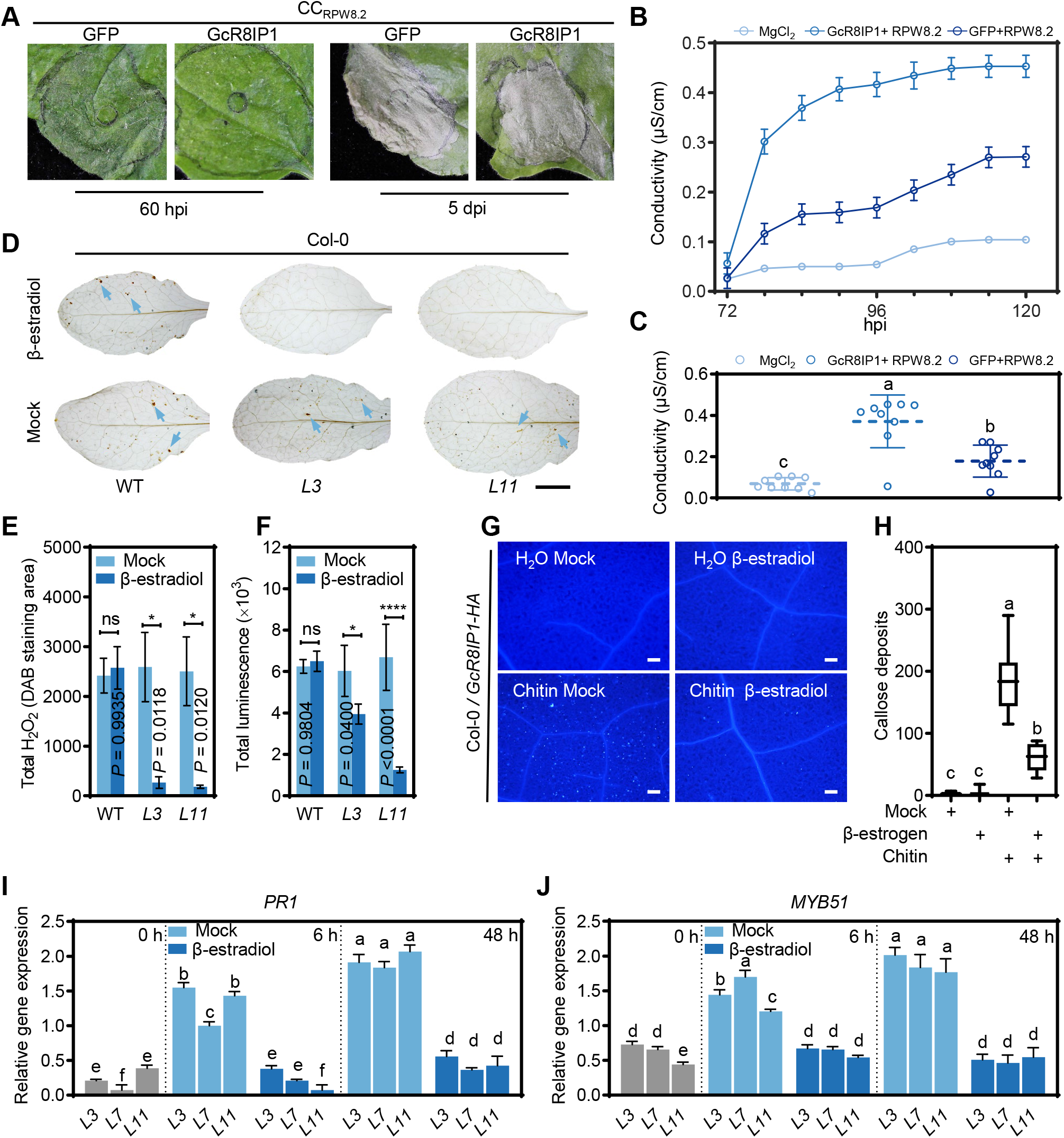
GcR8IP1 suppresses immune response in plants. A, Representative leaf sections show cell death phenotypes at the indicated time points. Leave of *N. benthamiana* were infiltrated with the agrobacterial strains containing CC_RPW8.2_ and GcR8IP1 or CC_RPW8.2_ and GFP. hpi and dpi indicate hours and days post infiltration, respectively. B, Electrolyte leakage analysis on the conductivity of the indicated leaf disks. RPW8.2 was co-expressed with GcR8IP1 or GFP in independent leaves. *N. benthamiana* leaves infiltrated with 1 mM MgCl_2_ were used as control. The conductivity was measured at the indicated time points after agrobacterium-infiltration. Error bars represent the mean ± s.d. (n = 3). C, Statistical analysis on electrolyte leakage from three biological repeats. The data are shown as mean ± s.d. (n = 9), and different letters indicate significant differences (*P* < 0.05) as determined by the one-way Tukey–Kramer test. D, Representative leaves after 3,3ʹ-diaminobenzidine (DAB) staining show H_2_O_2_ production in Col-0 (WT) and the indicated transgenic lines of XVE-*GcR8IP1*-*HA*-OE for β-estradiol inducible expression of GcR8IP1-HA in Col-0 background at 24 hours post inoculation of *G. cichoracearum* UCSC1. The leaves from the indicated lines were spray treated with β-estradiol or DMSO (mock) 12 h prior to the inoculation. Arrows indicate the position of representative DAB staining. Size bar, 1 cm. E, Quantification of the hydrogen peroxide (H_2_O_2_) production in the indicated lines from (D). ImageJ was used for quantification of DAB staining area. Error bars represent the mean ± s.d. (n = 3). Asterisks (*) and ns denote significant (*P* < 0.05) and no significant difference, respectively, determined by Student ’s *t*-test. F, Chemiluminescence assay shows elicitation of the oxidative burst induced by chitin in the indicated lines. Leaf disks punched from 4-week-old mature plants were treated with 100 µg/mL of chitin after 12 h spray-treated with β-estradiol or DMSO (mock). Signals were recorded for 30 min after treatment. Error bars represent the mean ± s.d. (n = 8). Asterisks (*, ****) denote significant differences (*P* < 0.05, *P* < 0.001, respectively; ns, no significant difference) as determined by Student ’s *t*-test. G, Representative images show callose deposition elicited by chitin in transgenic plants of XVE-*GcR8IP1*-*HA*-OE in Col-0 background after the indicated treatments. The rosette leaves were spray-treated with β-estradiol or DMSO (mock). After 12 h, the leaf discs were subjected to the treatment of 100 µg/mL of chitin or H_2_O. H, Quantification of callose depositions. ImageJ was used for quantification of callose deposition in (G). The data are shown as mean ± s.d. (n = 12), and different letters indicate significant differences (*P* < 0.05) as determined by the one-way Tukey–Kramer test. Similar results were observed in three independent experiments. I and J, Expression of the indicated defense-related marker genes in the indicated transgenic lines of XVE-*GcR8IP1*-*HA*-OE for β-estradiol inducible expression of GcR8IP1-HA in Col-0 background. The leaves from the indicated lines were spray-treated with β-estradiol or DMSO (mock) 12 h prior to the inoculation of *G. cichoracearum* UCSC1. Leaf samples were collected at the indicated time points. The data are shown as mean ± s.d. (n = 3), and different letters indicate significant differences (*P* < 0.05) as determined by the one-way Tukey–Kramer test.

To confirm GcR8IP1 suppressing plant immunity, we constructed *Arabidopsis* plants transgenic for *35Spro:GcR8IP1*-*GFP*, *35Spro:RFP*-*GcR8IP1* or *35Spro:GcR8IP1*-*RFP*. Unfortunately, we failed to detect GFP or RFP signals or proteins in the transgenic lines, although we detected the transcripts of *GcR8IP1*-*RFP* (Supplemental Figure S5A, B). Then, we exploited the XVE expression cassette to make the β-estradiol inducible expression of *GcR8IP1*-*HA* (XVE-*GcR8IP1*-*HA*-OE) in Arabidopsis accession Columbia-0 (Col-0 that lacks RPW8.1 and RPW8.2) or Warschau-1 (Wa-1 that contains RPW8.1 and RPW8.2 (Orgil et al., 2007)) following a previous report (Schlücking et al., 2013). GcR8IP1-HA was detected at 6, 12, and 24 hours after β-estradiol treatment and peaked at 12 hours in the transgenic lines in both Col-0 and Wa-1 backgrounds (Supplemental Figure S5C, D). These observations imply that GcR8IP1 only transiently accumulates when heterologously expressed in Arabidopsis.

After pathogen infection, less H_2_O_2_ accumulation was detected in the β-estradiol-treated plants of XVE-*GcR8IP1*-*HA*-OE transgenic lines in both Col-0 and Wa-1 background than in wild type (WT) and mock-treated transgenic lines at 24 h-post-inoculation of *Gc* UCSC1 (Figure 3D, E, Supplemental Figure S6A). The chitin-induced ROS accumulation was reduced by 40% to 50% in the XVE-*GcR8IP1*-*HA*-OE transgenic plants in comparison with that in the control plants (Figure 3F, Supplemental Figure S6B). Consistently, callose deposition was significantly less in the β-estradiol-treated Col-0/XVE-*GcR8IP1*-*HA*-OE plants than in the control plants upon chitin treatment (Figure 3G, H). The expression of two-defense-related marker genes, namely *PR1* and *MYB51*, was significantly less abundant in the estradiol-treated plants of Col-0/XVE-*GcR8IP1*-*HA*-OE than the mock plants at 6 hpi and 48 hpi (Figure 3I, J), indicating suppression. In contrast, although the expression of *PR1* and *MYB51* was significantly less in the estradiol-treated plants of Wa-1/XVE-*GcR8IP1*-*HA*-OE than the mock plants at 6 hpi, the expression was not different at 48 hpi (Supplemental Figure S6C, D), indicating delayed induction of these genes. These results suggest that GcR8IP1suppresses plant immunity, but such suppression is different between plants with or without the expression of RPW8.1 and RPW8.2.

### GcR8IP1 facilitates powdery mildew pathogenesis

To test the roles of *GcR8IP1* in PM pathogenesis, we inoculated the Col-0/XVE-*GcR8IP1*-*HA*-OE lines with *Gc* UCSC1 after β-estradiol treatment. Compared with the mock treatment, β-estradiol-treated plants exhibited enhanced susceptibility to PM (Figure 4A). The infection of PM leads to a decrease in photochemical efficiency that can be measured as chlorophyll fluorescence parameter Fv/Fm in the infected leaves (Kuckenberg et al., 2009). Indeed, chlorophyll fluorescence parameter Fv/Fm was significantly decreased in β-estradiol*-*treated Col-0/XVE-*GcR8IP1*-*HA*-OE lines in comparison to WT and mock-treated lines, implying colonized more PM (Figure 4B). Consistently, more conidiospores and fungal biomass were generated by β-estradiol-treated Col-0/XVE-*GcR8IP1*-*HA*-OE lines than by WT and mock-treated lines (Figure 4C, D), indicating enhanced susceptibility. These results indicate that ectopic expression of *GcR8IP1* results in enhanced susceptibility to PM in *Arabidopsis* likely via suppression of PTI.

**Figure 4.**
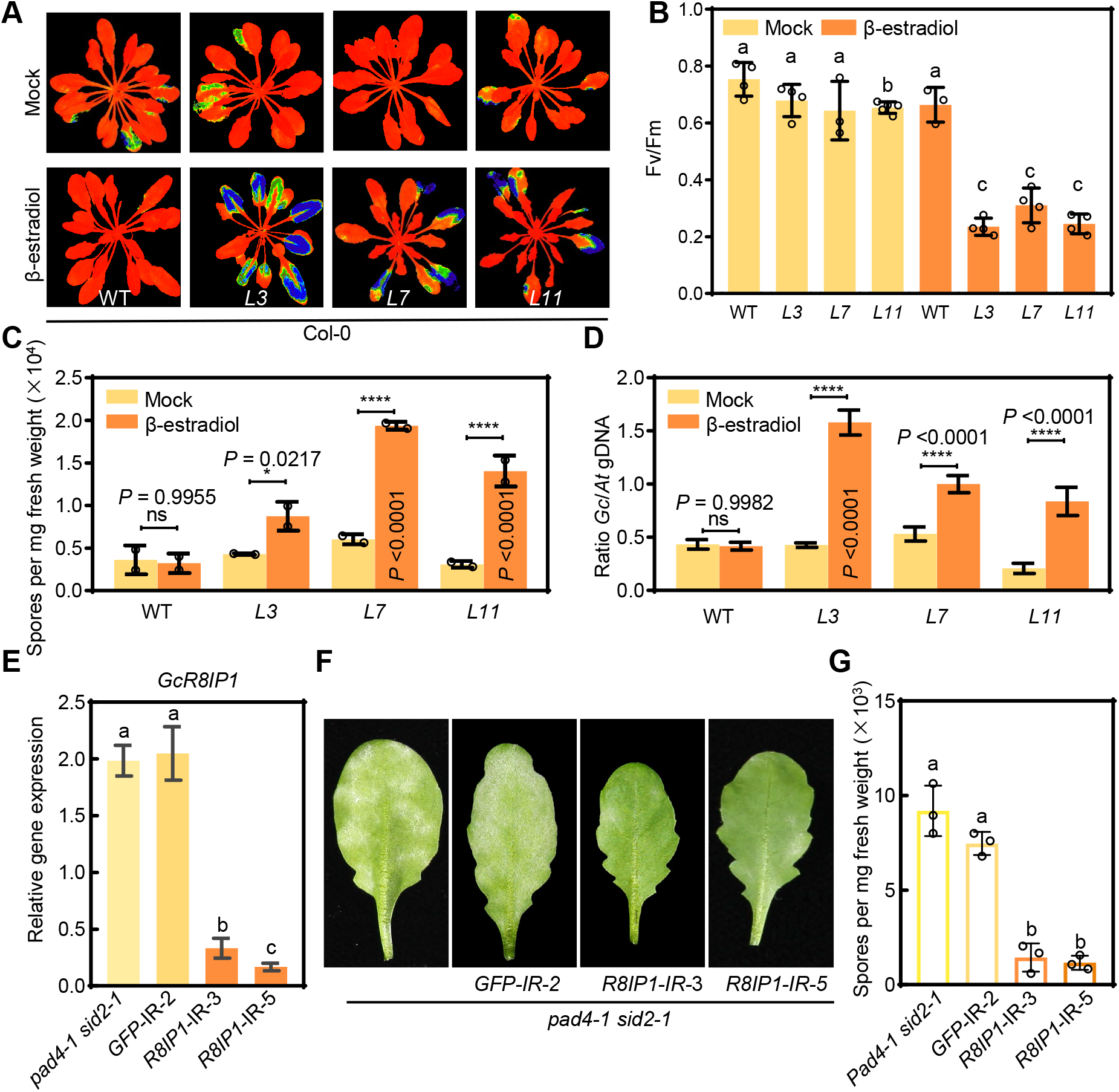
GcR8IP1 facilitates powdery mildew pathogenesis. A, Chlorophyll auto-fluorescence images indicate the severity of powdery mildew (PM) in the indicated transgenic lines of XVE-*GcR8IP1*-*HA*-OE for β-estradiol inducible expression of GcR8IP1-HA in Col-0 background in comparison with Col-0 (WT) at 10 days post inoculation (dpi) of *G*. *cichoracearum* UCSC1. The leaves from the indicated lines were spray-treated with β-estradiol or DMSO (mock) 12 h prior to the inoculation. B, Quantification analysis on the chlorophyll fluorescence parameter Fv/Fm from the indicated lines. The data are shown as mean ± s.d. (n = 5), and different letters indicate significant differences (*P* < 0.05) as determined by the one-way Tukey–Kramer test. C, Quantification analysis on the sporulation of PM from the indicated lines at 10 dpi. Error bars represent the mean ± s.d. (n = 6) and asterisks (*, ****) denote significant differences (*P* < 0.05, *P* < 0.001, respectively) as determined by Student ’s *t*-test. D, Quantification of relative PM biomass by qPCR in the indicated lines. Relative fungal biomass was calculated by the comparison between *G. cichoracearum GDSL-like lipase* gene and *A. thaliana Glyceraldehyde-3-phosphate dehydrogenase of plastid 2* gene (*GAPCP-2*) at 10 dpi of *Gc* UCSC1. The data are shown as mean ± s.d. (n = 3), and different letters indicate significant differences (*P* < 0.05) as determined by the one-way Tukey–Kramer test. E, Relative expression of *GcR8IP1* in PM-infected leaves from the indicated transgenic lines expressing interference small RNA for inducing silencing of *GFP* (GFP-IR) or *GcR8IP1* (R8IP1-IR) in the *pad4-1 sid2-1* background in comparison with *pad4-1 sid2-1* at 6 dpi of *Gc* UCSC1. The data are shown as mean ± s.d. (n = 3), and different letters indicate significant differences (*P* < 0.05) as determined by the one-way Tukey–Kramer test. F, PM disease phenotypes of the indicated lines. Photos were taken at 6 dpi of *Gc* UCSC1. G, Quantification analysis on the sporulation of PM from the indicated lines at 6 dpi. The data are shown as mean ± s.d. (n = 3), and different letters indicate significant differences (*P* < 0.05) as determined by the one-way Tukey–Kramer test. Similar results were obtained in three independent experiments.

To confirm the role of *GcR8IP1* in PM pathogenesis, we tried to reduce the expression of *GcR8IP1* in *Gc* UCSC1 during its growth in a susceptible *Arabidopsis* mutant *pad4-1 sid2-1* transgenic for *R8IP1*-*IR* and *GFP*-*IR* (Supplemental Figure S7), following a previous host-induced gene silencing (HIGS) approach (Nowara et al., 2010). Upon inoculation of *Gc* UCSC1 at 6 dpi, the expression of *GcR8IP1* was significantly down-regulated in the *R8IP1*-*IR* lines but not in the *GFP*-*IR* lines (Figure 4E), indicating effective HIGS of *GcR8IP1*. Disease assay indicated that *R8IP1-IR* lines exhibited less proliferation of PM than control and the *GFP-IR* lines at 6 dpi (Figure 4F). Consistently, conidiospore production was significantly decreased in *R8IP1*-*IR* lines in comparison with control and the *GFP-IR* lines (Figure 4G). These results indicate that HIGS of *GcR8IP1* results in compromised virulence of *Gc* UCSC1 on *pad4-1 sid2-1*. Thus, *GcR8IP1* is required for the full virulence of *Gc* UCSC1.

### Expression of GcR8IP1 increases the amount of nucleus-localized RPW8.2

To investigate the biological impact of GcR8IP1’s interaction with RPW8.2, we conducted a series of co-expression assays following a previous report (Huang et al., 2014). RPW8.2-GFP was localized mainly in the cytoplasm and rarely in the nucleus when *R82_pro_:RPW8.2-GFP* was co-expressed with *35S_pro_:2RFP-NLS* (Figure 5A). Intriguingly, RPW8.2-GFP was also in the nucleus when *R82_pro_:RPW8.2-GFP* was co-expressed with *35S_pro_:GcR8IP1-RFP* (Figure 5A, Supplemental Figure S8). The percentage of nuclei with RPW8.2-GFP signal was remarkably higher in the leaf epidermal cells co-expressing *R82_pro_:RPW8.2-GFP* with *35S_pro_:GcR8IP1-RFP* than those co-expressing *R82_pro_:RPW8.2-GFP* with *35Spro:2RFP-NLS* (Figure 5B). Consistently, RPW8.2-GFP was highly co-localized with GcR8IP1-RFP in the nucleus (Figure 5C-F). These data suggest that GcR8IP1’s interaction with RPW8.2 may facilitate nuclear localization of RPW8.2.

**Figure 5.**
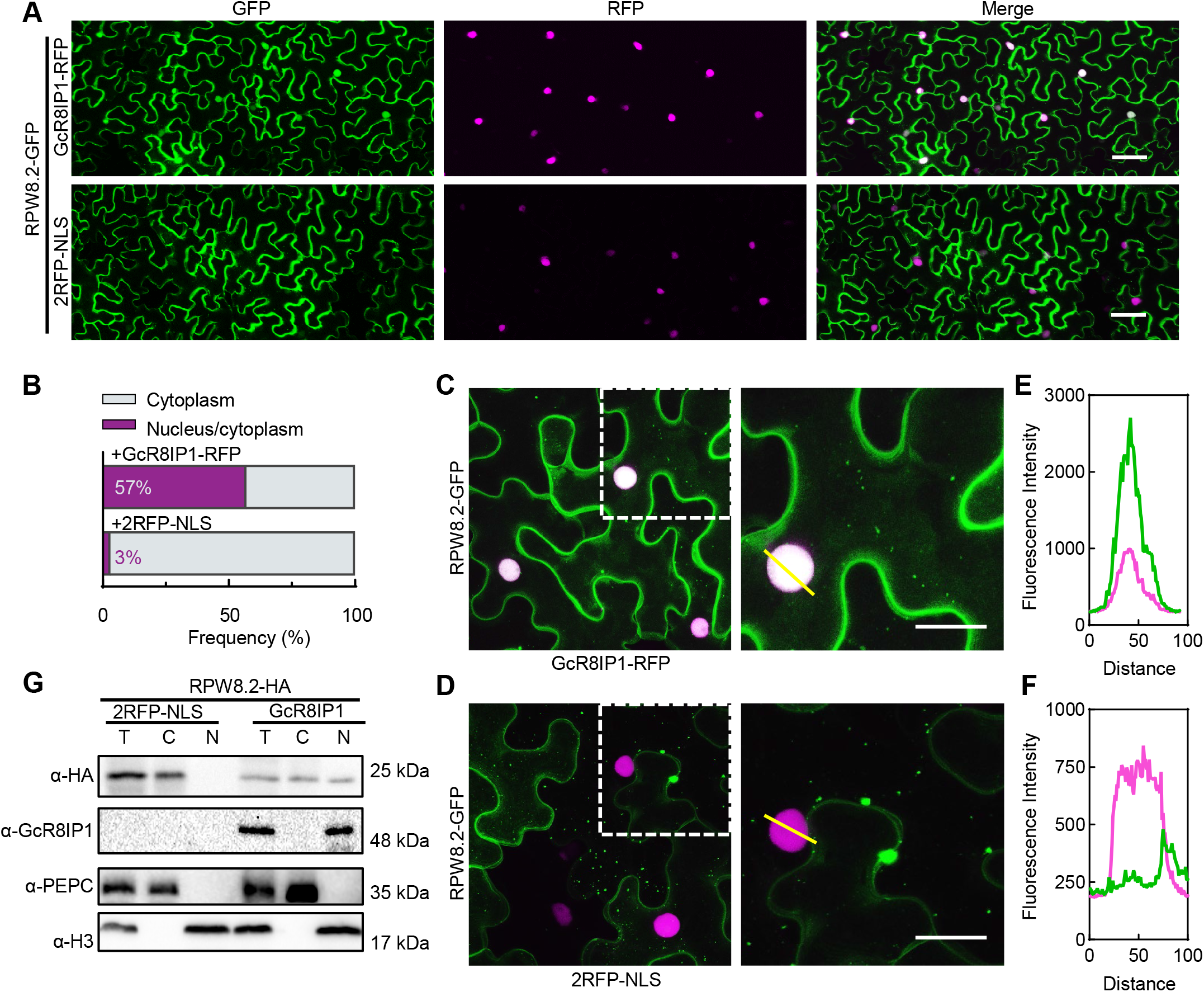
GcR8IP1 increases the partitioning of RPW8.2 in the nucleus. A, Representative confocal images show the subcellular localization of the indicated proteins. *R82_pro_:RPW8.2-GFP* was transiently co-expressed with *35S_pro_:GcR8IP1-RFP* (upper panel) and *35S_pro_:2RFP-NLS* (lower panel) in *N. benthamiana* leaves via Agrobacteria-mediated infiltration and images were acquired at 48 hours post infiltration. Size bars, 50 μm. B, Frequency of cells with RPW8.2-GFP in the nucleus. At least 200 cells were counted (refer to Supplemental Figure 8). C and D, Magnified confocal images show the subcellular localization of the RPW8.2-GFP when co-expressed with GcR8IP1-RFP (C) and 2RFP-NLS (D). Size bar, 20 μm. E and F, Scan line analysis on the fluorescence intensity of GFP and RFP at the position indicated by the lines in (C, D), respectively. G, Immunoblotting analysis on the partitioning of RPW8.2-HA in the nucleus and the cytoplasm when 35S_pro_:RPW8.2-HA was co-expressed with 35S_pro_:2RFP-NLS or 35S_pro_:GcR8IP1 in *N. benthamiana*. H3 and PEPC is a nuclear and a cytoplasmic marker, respectively.

Next, we prepared the cytoplasmic and nuclear fractions from total proteins extracted from the *N. benthamiana* leaves co-expressing *35S_pro_*:*RPW8.2-HA* with *35S_pro_:2RFP-NLS* or *35S_pro_:GcR8IP1* and examined the abundance of RPW8.2-HA by immunoblot analysis with two quality controls for the fractionation procedure, namely a cytosolic protein marker (phosphoenolpyruvate carboxylase [PEPC]) and a nuclear protein marker (histone H3). PEPC was detected only in the cytoplasmic fractions and histone H3 only in the nuclear fractions, indicating minimal contamination between the cytoplasmic and nuclear fractions (Figure 5G). Consistent with the subcellular localization, RPW8.2-HA was detected mainly in the cytoplasmic fractions but barely in the nuclear fractions when *35S_pro_:RPW8.2-HA* was co-expressed with *35S_pro_:2RFP-NLS*. In contrast, RPW8.2-HA was detected both in the cytoplasmic and the nuclear fractions when *35S_pro_:RPW8.2-HA* was co-expressed with *35S_pro_:GcR8IP1* (Figure 5G), indicating that expression of GcR8IP1 alters the nucleocytoplasmic partitioning of RPW8.2.

### Nucleus-localized RPW8.2 promotes the activity of the *RPW8.2* promoter

*RPW8.2* is induced by PM infection presumably via SA-dependent feedback amplification (Xiao et al., 2003; Wang et al., 2009). To test whether the nucleus-localized RPW8.2 is crucial for such a feedback amplification, we measured the expression of a firefly luciferase (LUC) reporter under control of the *RPW8.2* promoter (*R82_pro_:LUC*) when it was co-expressed with a wild type RPW8.2 (*35S_pro_:RPW8.2*), a nuclear location signal (NLS)-fused RPW8.2 (*35S_pro_:RPW8.2-HA-NLS*), a nuclear export signal (NES)-fused RPW8.2 (*35S_pro_:RPW8.2-HA-NES_PKI_*) following a previous report (Huang et al., 2014), or a mutated NLS-fused RPW8.2 (*35S_pro_:RPW8.2-HA-nls*) (Huang et al., 2019). The activity of *R82_pro_:LUC* was enhanced when it was co-expressed with *35S_pro_:RPW8.2-HA-NLS*, but suppressed when co-expressed with *35S_pro_:RPW8.2-HA-NES_PKI_*, compared with the co-expression with the empty vector (EV), the wild-type control, and *35S_pro_:RPW8.2-HA-nls* (Fig 6a), implying that nucleus-localized RPW8.2 increases the activity of the *RPW8.2* promoter.

**Figure 6.**
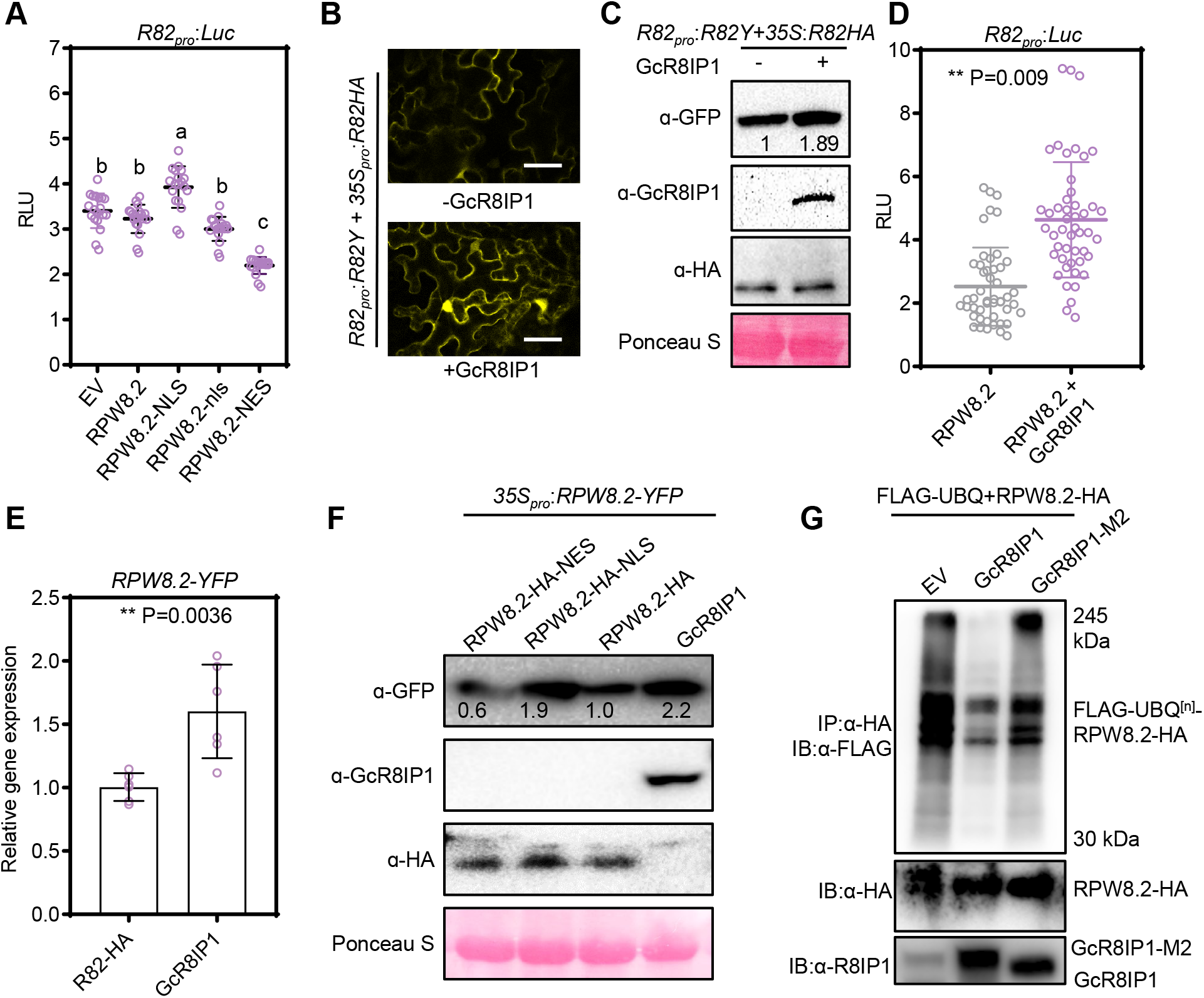
GcR8IP1 increases RPW8.2 accumulation. A, Dual-luciferase assay shows the relative luciferase activities between firefly luciferase and renilla luciferase. Firefly luciferase was expressed from the *RPW8.2* promoter (*R82_pro_:Luc*) and co-expressed with empty vector (EV), *RPW8.2*, *RPW8.2-NLS*, RPW8.2-nls and RPW8.2-NES, respectively, which were driven by the *35S* promoter. 35S-expressed *Renilla luciferase* was used as the internal reference. The results are presented as relative luminescence units (RLU) normalized to the ratio of firefly luciferase/renilla luciferase. The data are shown as mean ± s.d. (n = 40), and different letters indicate significant differences (*P* < 0.05) as determined by the one-way Tukey–Kramer test. B, Representative confocal images of RPW8.2-YFP (*R82_pro_:R82Y*) co-expressed with RPW8.2-HA (35S_pro_:R82HA) plus / minus GcR8IP1 in *N. benthamiana*. C, Immunoblotting analysis of (B) shows the accumulation of RPW8.2-YFP. Numbers below the blot indicate relative abundances of RPW8.2-YFP. D, Dual-luciferase assay shows the relative luciferase activities between firefly luciferase and renilla luciferase. Firefly luciferase was expressed from the *RPW8.2* promoter (*R82_pro_:Luc*) and co-expressed with RPW8.2 or RPW8.2 plus GcR8IP1. *Renilla luciferase* expressed from the 35S promoter was used as internal reference. Error bars represent the mean ± s.d. (n = 48) and asterisks (**) denote significant differences (*P* < 0.01) as determined by Student ’s *t*-test. E, Quantitative PCR assay shows the relative transcript abundance of *RPW8.2-YFP*. RPW8.2-YFP from its native promoter was transiently co-expressed with RPW8.2-HA from the *35S* promoter (R82-HA) or GcR8IP1 in *N. benthamiana*. Error bars represent the mean ± s.d. (n = 8) and asterisks (**) denote significant differences (*P* < 0.01) as determined by Student ’s *t*-test. F, Immunoblotting analysis on the abundance of RPW8.2-YFP when it was co-expressed with the indicated proteins. Numbers below the blot indicate relative abundances of RPW8.2-YFP determined by optical density quantitative analysis on the indicated co-expressions. G, Immunoblotting analysis on the ubiquitination of RPW8.2-HA. Total proteins were extracted from *N. benthamiana* leaves co-expressing FLAG-UBQ and RPW8.2-HA with EV, FLAG-GcR8IP1 and FLAG-GcR8IP1-M2, respectively. Protein complexes were pulled down using anti-HA agarose beads and the co-precipitation of FLAG-UBQ-tagged-RPW8.2-HA (FLAG-UBQ^[n]^-RPW8.2-HA) was detected by anti-FLAG antibody. The RPW8.2-HA was detected by anti-HA antibody. The GcR8IP1 and GcR8IP1-M2 were detected by anti-R8IP1 antibody.

To evaluate the effect of the interaction between RPW8.2 and GcR8IP1 on the *RPW8.2* promoter activity, we examined the amount of RPW8.2-YFP expressed from the *RPW8.2* promoter (*R82_pro_:R82Y*) when it was co-expressed with RPW8.2-HA (*35S_pro_:R82HA*), or with RPW8.2-HA plus GcR8IP1 (*35S_pro_:GcR8IP1*) in *N. benthamiana*. The RPW8.2-YFP signal from *R82_pro_:R82Y* was obviously more intense when *R82_pro_:R82Y* was co-expressed with *35Spro:RPW8.2-HA* plus *35S_pro_:GcR8IP1* than co-expressed with *35S_pro_:R82HA* (Figure 6B). Immune blotting analysis confirmed that more RPW8.2-YFP accumulated when *R82pro:R82Y* was co-expressed with *35S_pro_:R82HA* plus *35S_pro_:GcR8IP1* than co-expressed with *35S_pro_:R82HA* (Figure 6C), indicating that the co-expression of GcR8IP1 may enhance the transcription of *R82_pro_:R82Y* or stabilize RPW8.2-YFP, or both. Consistently, the activity of *R82_pro_:LUC* was significantly increased in its co-expression with *35S_pro_:RPW8.2* plus *35S_pro_:GcR8IP1* than its co-expression with *35S_pro_:RPW8.2* (Figure 6D), indicating that co-expression of GcR8IP1 enhances the *RPW8.2* promoter activity presumably due to the increase of the partitioning of RPW8.2 in the nucleus.

To confirm the effect of the partitioning of RPW8.2 on the activity of the *RPW8.2* promoter, we compared the transcript abundance of *RPW8.2-YFP* in leaves co-expressing *R82_pro_:RPW8.2-YFP* with *35S_pro_:R82HA* or *35S_pro_:GcR8IP1*. The transcripts of *RPW8.2-YFP* were more abundant when *R82_pro_:RPW8.2-YFP* was co-expressed with *35S_pro_:GcR8IP1* than when it was co-expressed with *35S_pro_:RPW8.2-HA*, implying that nuclear-localization of RPW8.2 promotes the activity of the *RPW8.2* promoter (Figure 6E).

To test whether GcR8IP1 also stabilizes RPW8.2, we examined the accumulation of RPW8.2-YFP from co-expression of *35S_pro_:RPW8.2-YFP* with *35S_pro_:GcR8IP1*, *35S_pro_:RPW8.2-HA-NLS*, *35S_pro_:RPW8.2-HA-NES_PKI_*, or *35S_pro_:RPW8.2-HA*. Again, the most abundant accumulation of RPW8.2-YFP was detected when *35S_pro_:RPW8.2-YFP* was co-expressed with *35S_pro_:GcR8IP1*, followed sequentially by when it was co-expressed with *35S_pro_:RPW8.2-HA-NLS*, *35S_pro_:RPW8.2-HA*, and *35S_pro_:RPW8.2-HA-NES_PKI_* (Figure 6F), indicating that GcR8IP1 also prevents RPW8.2 from degradation. Previously, RPW8.2 was reported to be degraded via 26S proteasome-mediated pathway and vacuole-dependent pathway (Huang et al., 2019). Consistently, Ubiquitination (UBQ) assay detected much less ubiquitinated RPW8.2-HA when *35S_pro_:RPW8.2-HA* was co-expressed with *35S_pro_:GcR8IP1* than its expression alone or co-expressed with *35S_pro_:GcR8IP1-M2* (Figure 6G), indicating that co-expression of GcR8IP1 reduces RPW8.2 ubiquitination.

### RPW8.2 and GcR8IP1 are engaged in a molecular warfare at the infection sites

To test whether GcR8IP1 triggers RPW8.2-mediated immunity against PM as the current concept of ETI, we first examined reactive oxygen species (ROS) production induced by the expression of RPW8.2 in *N*. *benthamiana*. To our surprise, ROS production was significantly suppressed when *35S_pro_:RPW8.2* was co-expressed with *35S_pro_:GcR8IP1* (Supplemental Figure S9). We then exploited the transgenic lines with β-estradiol inducible expression of GcR8IP1-HA in Wa-1 background (Wa-1/XVE-*GcR8IP1*-*HA*-OE). Both β-estradiol- and mock-treated Wa-1/XVE-*GcR8IP1*-*HA*-OE plants displayed disease resistance phenotypes similar to Wa-1 (Figure 7A). However, the transgenic lines of β-estradiol treatment supported significantly more fungal biomass than Wa-1 and mock treatment (Figure 7B). Consistently, photochemical efficiency assay showed that relative chlorophyll fluorescence (Fv/Fm) was decreased in β-estradiol treated Wa-1/XVE-*GcR8IP1*-*HA*-OE lines, indicating more PM fungal growth on the GcR8IP1-expressed leaves (Supplemental Figure S10A, B). Nevertheless, the β-estradiol treated Wa-1/XVE-*GcR8IP1*-*HA*-OE plants still produced comparable counts of conidiospores as the control plants, indicating effective resistance (Figure 7C).

**Figure 7.**
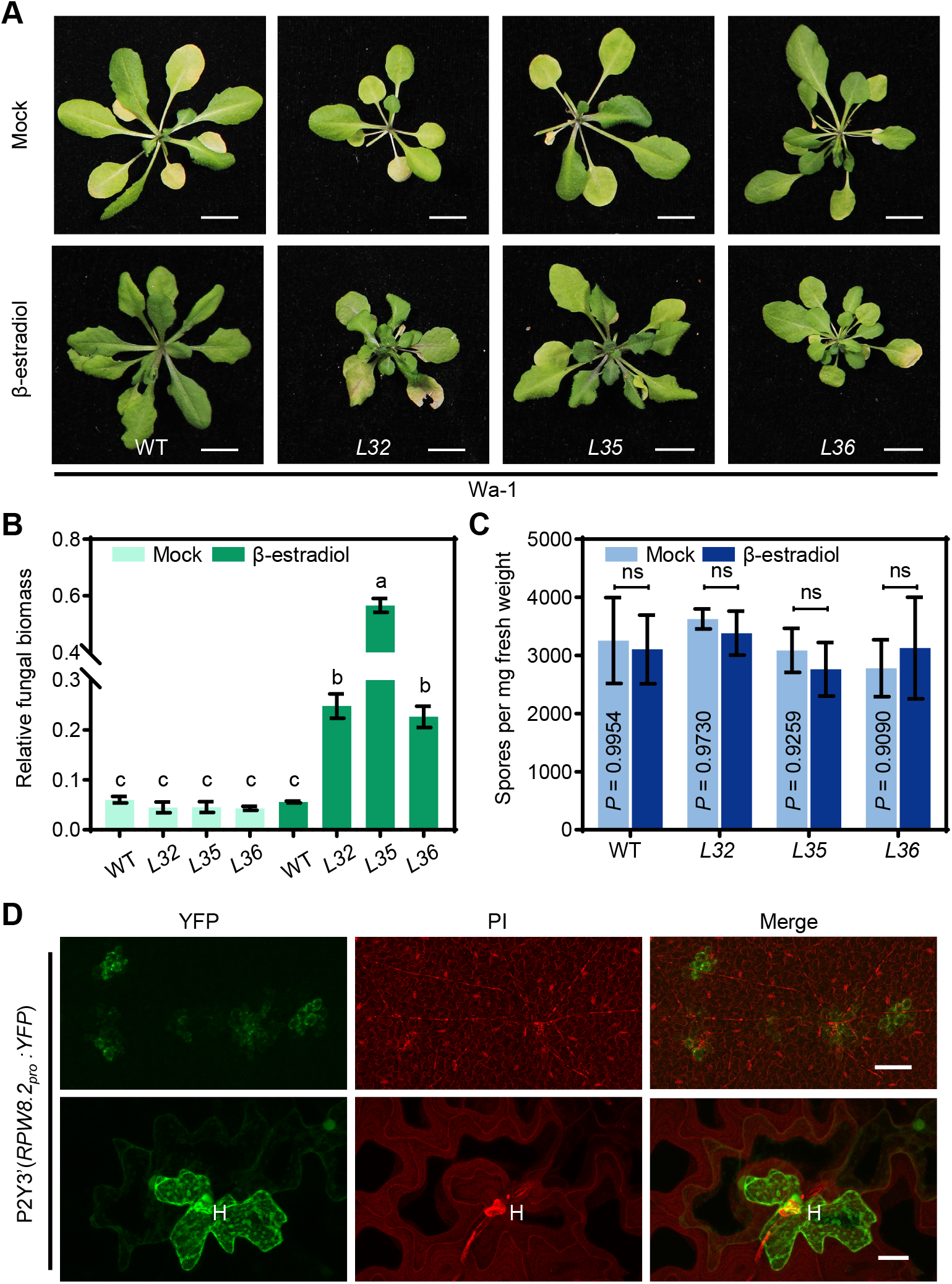
The molecular warfare engaged by GcR8IP1 and RPW8.2 at the infection site. A, Representative plants show the resistance phenotype of the indicated transgenic lines of XVE-*GcR8IP1*-HA-OE for β-estradiol inducible expression of *GcR8IP1-HA* in the Wa-1 (WT) background at 10 days post inoculation of *G. cichoracearum* UCSC1. The indicated lines were spray-treated with β-estradiol or DMSO (mock) 12 h prior to the inoculation. B, Quantification of powdery mildew (PM) biomass by qPCR in the indicated lines. Relative fungal biomass was calculated by the comparison between *G. cichoracearum GDSL-like lipase* gene and *A. thaliana Glyceraldehyde-3-phosphate dehydrogenase of plastid 2* gene (*GAPCP-2*) at 10 dpi of *G. cichoracearum* UCSC1. The data are shown as mean ± s.d. (n = 3), and different letters indicate significant differences (*P* < 0.05) as determined by the one-way Tukey–Kramer test. C, Quantification analysis on the sporulation of PM from the indicated lines at 10 dpi. The data are shown as mean ± s.d. (n = 3), and ns indicates no differences as determined by Student ’s *t*-test. Similar results were obtained in three independent experiments. D, Confocal images show the expression of YFP from the *RPW8.2* promoter in the transgenic plants P2Y3’ at 2 days post inoculation of *G. cichoracearum* UCSC1. H, haustorium. PI, propidium iodide. Size bars, 100 μm (upper) and 20 μm (lower).

The transcription of *RPW8* was positively regulated via an SA-dependent feedback loop and *RPW8.2* expression continually increases from 1 dpi with *Gc* UCSC1 and remained at high abundance at 7 dpi (Xiao et al., 2003, 2005). Consistently, the expression of *RPW8.2-YFP* from the *RPW8.2* promoter was highly up-regulated at 24 h, reaching approximately 1269 times higher than that at 0 h upon the treatment of the SA-functional analogue benzo (1,2,3) thiadiazole-7-carbothioic acid (BTH) (Supplemental Figure S11A). Surprisingly, the accumulation of RPW8.2-YFP protein was still undetectable at this time point (Supplemental Figure S11B), indicating that RPW8.2 in uninfected cells (in the absence of GcR8IP1) is constantly removed, which is consistent with our earlier report that RPW8.2 is turned over via both the 26s proteasome and the vacuole-dependent pathway in the cytoplasm (Huang et al., 2019). However, strong YFP expression from the *RPW8.2* promoter was detectable in haustorium-infected epidermal cells while only weak YFP signal was occasionally observed in adjacent cells of P2Y3’ transgenic lines expressing YFP from the *RPW8.2* promoter (Figure 7D). These data suggest that RPW8.2 in haustorium-invaded cells undergoes a rapid and strong self-amplification while RPW8.2 in uninfected cells is rapidly removed via the 26S proteasome and vacuole-dependent degradation, hence supporting a role of GcR8IP1 in inducing RPW8.2 expression and inhibiting RPW8.2’s degradation, which lead to a strong and sustained up regulation of RPW8.2 at the infection sites.

## Discussion

During canonical ETI, plant NLR receptors detect their cognate effectors, i.e., avirulence (Avr) factors to initiate defense responses summited by the hypersensitive response (HR) (Jones and Dangl, 2006). Here, we found that the PM effector GcR8IP1 acts like an Avr that activates RPW8.2. GcR8IP1 physically associates with RPW8.2 to increases its partitioning into the nucleus, which in turn, amplifies *RPW8.2* expression to boost immunity.

GcR8IP1 is a secreted effector delivered into the nuclei of host cells (Figure 2C). Even though no canonical SP was predicted at the N-terminus of GcR8IP1 by online prediction software, the first 16 to 40 aa achieves secretion of a reporter peptide in yeast or in planta (Figure 2A, 2B), indicating a SP different from the known five types of SPs (Teufel et al., 2022). Moreover, immunolocalization assays demonstrated that GcR8IP1 forms puncta in the nucleus of haustorium-invaded epidermal cells of Arabidopsis plants upon inoculation of *Gc* UCSC1. Such GcR8IP1-positive puncta are similar to those detected in BiFC that indicates direct interaction between GcR8IP1 and RPW8.2 (Figure 1F and Figure 2C). In contrast, GcR8IP1-GFP was detected as diffuse signal in the nucleus when transiently expressed in *N*. *benthamiana* (Supplemental Figure S4). These data indicate that GcR8IP1 is secreted by *Gc* UCSC1 and delivered into host cells and imply that its interaction with RPW8.2 or homologs in Arabidopsis may be attributable to GcR8IP1’s puncta localization in the nucleus. We also attempted to verify the secretion of GcR81IP1 in a model fungus *M. oryzae* but failed to detect GcR8IP1-GFP in transgenic *M. oryzae* strain. This could be due to rapid protein turnover when GcR8IP1 is heterologously expressed in *M. oryzae*. This could also explain why GcR8IP1 was not detectable when ectopically expressed in Arabidopsis from the *35S* promoter (Supplemental Figure S5). Thus, GcR8IP1 might be under tight post translational regulation in *Gc* UCSC1. Moreover, we also found that *GcR8IP1* exhibits increased in planta expression at 3-4 dpi and 6-7 dpi when haustorial establishment and sporulation occur, respectively, during infection (Supplemental Figure S3). Ectopic expression of GcR8IP1 in *N*. *benthamiana* and Arabidopsis suppressed plant immunity and facilitated PM infection (Figure 3 and Supplemental Figure S6, 9-10). HIGS of GcR8IP1 significantly impaired *Gc* UCSC1’s infectivity (Figure 4). Therefore, GcR8IP1 acts as a secreted effector required for pathogenicity of PM.

PM infection remarkably induces the expression of RPW8.2, which is partitioned to the EHM, the nucleus and the cytoplasm (Wang et al., 2009; Huang et al., 2019). The EHM-targeting of RPW8.2 is obviously induced upon the establishment of the haustorial complex. Here, we found that PM infection also impacts the partitioning of RPW8.2 into the nucleus via its association with GcR8IP1. Apparently, when GcR8IP1 is delivered to the nucleus of the host cell upon PM infection, its association with RPW8.2 facilitates the nuclear localization of RPW8.2, thus increasing the partitioning of RPW8.2 in the nucleus (Figure 5). Because RPW8.2 is known to engage an SA-signaling–dependent transcriptional self-amplification circuit (Xiao et al., 2003, 2005), increased nucleus-localized RPW8.2 further activates the *RPW8.2* promoter, leading to a rapid transcriptional self-amplification (Figure 6A, 6D, 6F, and 8B). Intriguingly, our data also suggest that GcR8IP1 decreases the ubiquitination of RPW8.2 (Figure 6G) via an unknown mechanism. One possibility is that nucleus-localized RPW8.2 may be protected from ubiquitination. Thus, the combinatorial impacts of GcR8IP1 on RPW8.2’s expression and accumulation provide a detailed mechanistic explanation as to why RPW8.2 is highly and specifically induced and functions in haustorium-invaded cells upon PM infection (Figure 8).

**Figure 8.**
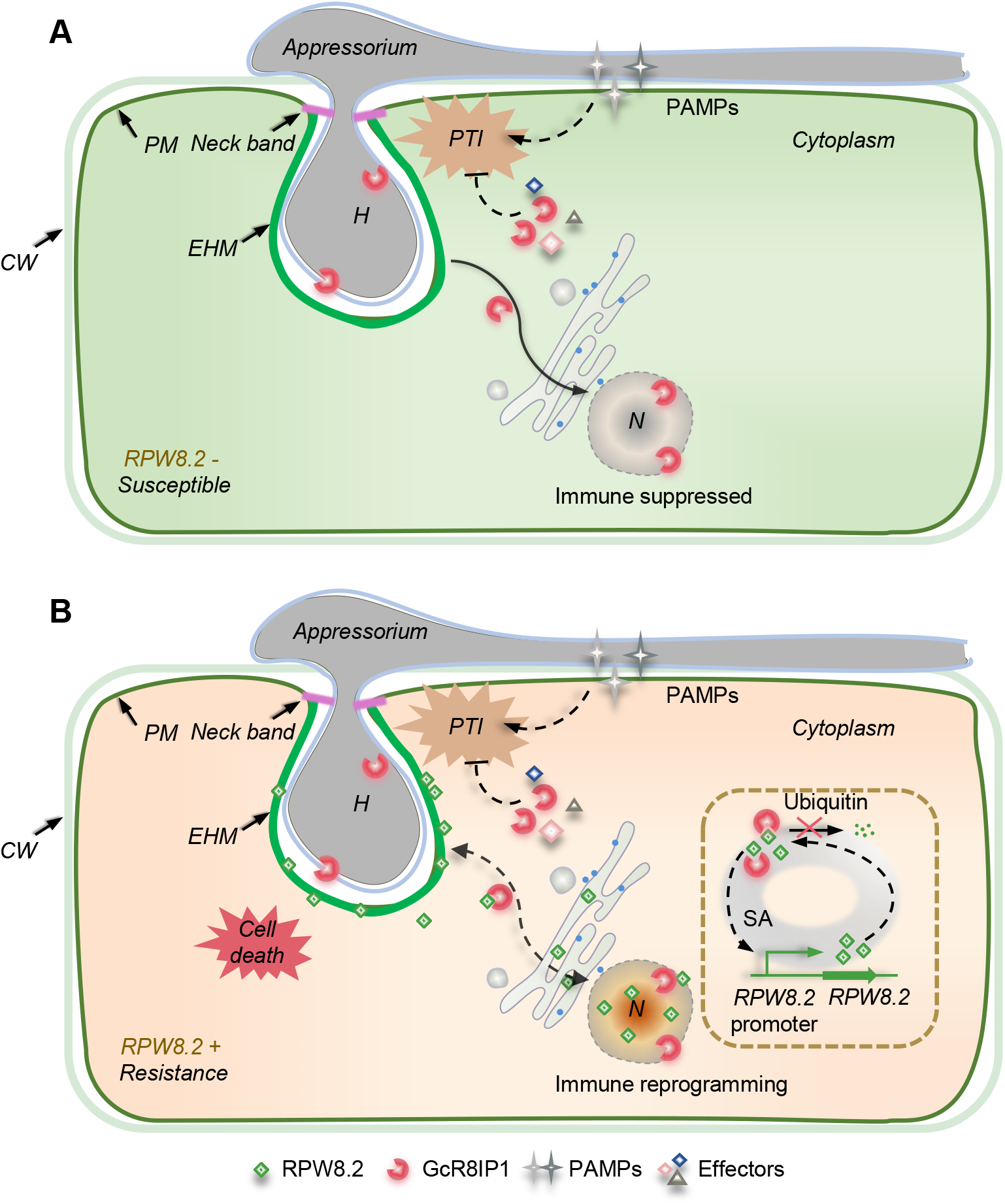
A working model for GcR8IP1-induced transcriptional amplification of RPW8.2 to activate immunity against powdery mildew. A, In the plants without RPW8.2 expression, PM delivers GcR8IP1 into the host to suppress immunity, presumably via suppression of PTI. B, In the plants with RPW8.2 expression, GcR8IP1 stabilizes RPW8.2 via suppressing its ubiquitination and increases its nuclear partitioning. Nucleus-localized RPW8.2 activates defense and the increased nucleus-localization further increases its expression via a SA-dependent transcriptional amplification, leading to broad-spectrum resistance to powdery mildew. CW, cell wall. EHM, extrahaustorial membrane. H, haustoria. PM, plasma membrane.

*GcR8IP1* is a highly conserved gene found in all sequenced powdery mildew genomes, suggesting that GcR8IP1 is a key virulence factor of PM fungi (Supplemental Figure S1). Because GcR8IP1 is conserved in PM fungi, RPW8.2 may bind homologs of GcR8IP1 from other PM fungi. Thus, the interaction of RPW8.2 with GcR8IP1 may in part explain why RPW8.2 confers broad-spectrum resistance to all infectious PM fungi tested (Xiao et al., 2001). However, the role of GcR8IP1-RPW8.2 interaction in immunity is mechanistically distinct from that of the conventional Avr-R recognition in ETI against PM, such as the MLA-AVR_A_ recognition in barley-PM and Pms-AvrPms in wheat-PM interactions (Saur et al., 2019; Bourras et al., 2019; Bettgenhaeuser et al., 2021). First, RPW8.2 may function as an executor rather than an immune receptor to recognize GcR8IP1 and initiate defense signaling. RPW8.2 seems to be passively hijacked by GcR8IP1 to localize to the nucleus where its accumulation triggers a SA-dependent self-amplification, leading to defense activation (Figure 8). Hence, it is possible that RPW8.2 may act as a decoy and the real host target of GcR8IP1 might be the RPW8-like domain of the CC_R_-NLRs. Second, the molecular warfare during canonical ETI is featured with a rapid HR within hours in the cases of bacteria, or 2 days with fungi and the interacted R-Avr triggers cell death when they are transiently co-expressed (Saur et al., 2019; Bourras et al., 2019; Bettgenhaeuser et al., 2021; Sun et al., 2021), whereas PM-induced RPW8-mediated HR occurs at 3 dpi or later (Xiao et al., 2003; Wang et al., 2009). This is consistent with our data that GcR8IP1 obviously suppresses RPW8.2-mediated immunity in both transient and stable expression assays in early infection stages when RPW8.2 protein expression level is rather low (Figure 3A-C, Figure 7B, Supplemental Figure S6). However, when RPW8.2 gets amplified above a threshold level as a consequence of its increased nuclear localization and accumulation due to its interaction with GcR8IP1, RPW8.2-mediated defense does not only offset the suppression of immunity by GcR8IP1, but also greatly stimulates EDS1- and SA-dependent signaling, leading to HR and restriction of fungal sporulation at day 3 and afterward (Figure 3A, 7A and 7C). Thus, the molecular interplay between GcR8IP1 and RPW8.2 is quite complicated, and its outcome depends on the basal level of SA-dependent defense, which determines the initial expression level of RPW8.2, and the level of GcR8IP1, which is probably determined by the quantity and activity of PM haustoria in the invaded host cells. Such a complex interplay offers a plausible new explanation for PM-induced and RPW8.2 dosage-dependent broad-spectrum resistance against PM fungi (Figure 8B).

## Materials and methods

### Plant Materials and Growth conditions

*Arabidopsis thaliana* accessions Wa-1, Col-0 and *pad4-1 sid2-1* double mutant (Wen et al., 2011) were used for transgenic analysis. Seeds were sown directly on soil, and vernalized at 4 °C for 2 days before moving to a growth room with 10 hours of light and 14 hours of dark, 75% humidity under 21 °C. *Nicotiana benthamiana* was always cultivated in the same conditions. The floral dip was used for plant transformation as described (Clough and Bent, 1998). Transgenic plants were selected in soil by spraying 15 mg/L Basta.

### Yeast Two-Hybrid Assay

Y2H assays were performed according to the manufacturer’s protocol (Clontech). For Y2H screening, full-length coding sequences of *RPW8.2* were fused to the MCS in the pGBKT7 vector for bait construction and the screening of an Arabidopsis cDNA fusion library prepared with mRNA extracted from *Gc* UCSC1 infected leaves.

### *Agrobacterium*-Mediated Transient Expression Assay and Protein Subcellular Localization Analysis

Constructs harboring different genes were transformed individually into *Agrobacterium tumefaciens* strain GV3101. *Agrobacterium* cultures were harvested and resuspended in infiltration buffer (1mM MgCl_2_, 10mM MES pH=5.7, and 200 μM acetosyringone) to OD_600_ = 0.8 for infiltration (Wydro et al., 2006). In co-expression assays, equal volume of the indicated cultures was mixed thoroughly for infiltration. The plants were then maintained in regular growth conditions for 48 h before microscopic observation or immunoblotting. The fluorescence signal of GFP or RFP was observed using a confocal laser scanning microscope.

### Protein Expression and Purification from *Escherichia coli*

The pGEX6p-1 constructs transfected into *Escherichia coli* strain Rosetta (DE3). The expression and purification of recombinant proteins in the *E. coli* were performed according to the manual provided by GE healthcare. Briefly, *E. coli* was first grown in liquid Luria-Bertani medium to reach OD_600_ = 0.3, and then supplied with 20 mM Isopropyl β-D-1-Thiogalactopyranoside for further grown until OD_600_ = 2.0. The culture was harvested and the recombinant protein purified using glutathione resin according to the manufacturer’s instructions (GE healthcare, 17075601).

### In Vitro Protein–Protein Interaction Assay

The *in vitro* protein-protein interaction assay was performed as described (Chang et al., 2013) with modifications. Briefly, 1 mg purified GST and GST fusion protein was incubated with 50 μL glutathione resin 4 h at 4°C. Then the resin washed with buffer A (150 mM NaCl, 50 mM Tris-Cl, pH 7.4, 0.1% Tween 20, protease inhibitor cocktail, and 1mM DTT) and incubated 2 hours with 5 mL protein isolated from 2.0 g *N. benthamiana* leaves transient-expressing interest protein. After incubation, the beads were washed 6 times with buffer A, resuspended in 4× SDS sample buffer and boiled for 5 min. The resulting proteins were separated on SDS-PAGE gels and examined using an α-GST and α-HA antibody, respectively.

### Co-IP and immunoblotting analyses

In Co-IP assays, RPW8.2-FLAG transiently co-expressed with GcR8IP1-HA or GcR8IP1-M2-HA, respectively in *N. benthamiana*. Total protein was extracted by extraction buffer (50 mM HEPES, pH 7.5, 150 mM KCl, 1 mM EDTA, 0.5% Triton X-100, 1 mM DTT, and 1× proteinase inhibitor cocktail). For α-HA IP, the total protein was incubated with 50 µL agarose-conjugated α-HA antibody for 4 h and then washed 8 times with the extraction buffer. The total protein was eluted with 0.5 mg/mL 3× HA peptide for 0.5 h and then separated by SDS-PAGE and detected by α-HA and α-FLAG immunoblot. Cytoplasmic and nuclear protein fractionation was performed as described (Wang et al., 2011). The resulting proteins were separated on SDS-PAGE gels and examined using an α-R8IP1, α-HA, α-H3 and α-PEPC antibody, respectively.

### Luciferase Complementation Imaging Assay

For LCI assay, *N. benthamiana* leaves were co-infiltrated with the agrobacteria culture carrying the nLuc and cLuc derivative constructs. The infiltrated leaves were sprayed with 1 mM luciferin at 48 h after infiltration. The images were taken by cooled charge coupled device imaging apparatus and quantitative LUC activity was measured by a luminometer.

### Immunofluorescence assay

The leaves were detached from Arabidopsis rosette at 3 d post *Gc* UCSC1 inoculation and treated with the following steps including deparaffinization in xylene, rehydration in a graded ethanol series and air-drying. The treated leaves were chipped into slices with 6-8 μm thickness, then washed by phosphate-buffered saline with 0.1% Triton X-100, pH7.4 (PBS-T). The slices were immersed 30 min in blocking buffer (10% normal goat serum in PBS-T and then transferred into incubation buffer (1% normal goat serum in PBS-T) that contains the α-R8IP1 with 1: 300 dilution for 4 h incubation. The slices were washed by PBS-T for 3× 5 min and incubated for 1 h with a 1: 600 dilution of the secondary antibody (goat anti-rabbit Alexa Fluor 488, Thermo). The slices were washed with the PBS-T buffer for three times again. Finally, the slices were placed on slides with the PBS-glycerol (1:1) solution and carefully overlaid with a glass coverslip for observation.

### Measurement of the maximum photochemical efficiency of PSII

The 8-week-old Arabidopsis were used to determine the maximum photochemical efficiency of PSII at 10 dpi after *Gc* UCSC1 inoculation. The potential quantum efficiency of PSII was measured by calculating the ratio Fv/Fm [(Fm–Fo)/Fm] (Butler and Kitajima, 1975).

### Diaminobenzidine (DAB) staining assay

3,3’-diaminobenzidine (DAB) was used to stain *in situ* H_2_O_2_ in the leaves. 24 h after inoculation The leaves infected with PM were detacted at 24 hpi and immersed in 1 mg/mL 3,3’-diaminobenzidine solution (0.2M sodium phosphate buffer, pH 7.0) overnight in dark and then destained by 95% ethanol. The images were taken under microscope to record DAB stains.

### ROS assay

Production of ROS was assayed as described (Li et al., 2010). Briefly, 3-mm leaf discs were detached and soaked in ddH_2_O in the wells of opacity 96-well plates for 8 h. The ddH_2_O was carefully removed before adding 200 mL buffer containing 20 mM luminol and 1 mg horseradish peroxidase with 100 μg/mL chitin. Luminescence was determined by a luminometer.

### Host-Induced Gene Silencing Assay

The fragment of *GcR8IP1* covering the RING domain-encoding sequence from 27 bp to 382 bp was used for silencing *GcR8IP1* via HIGS. The fragment of GFP from 62 bp to 462 bp was used as a negative control in this assay. The fragments were cloned into pKANNIBAL vector respectively. The primers used for generate invented repeat constructs are listed in Supplemental Table 1. Four-week-old transgenic lines were inoculated with *Gc* UCSC1.

### RT-PCR and Real-Time PCR Analysis

For gene transcription analysis, total RNA was extracted from *Arabidopsis* or *N. benthamiana* leaves using TRizol-Reagent (Invitrogen 15596018) and then treated with RNase-free DNase I (Takara 2270B) to remove the potential DNA contamination. The first-strand cDNA was synthesized from 1μg of total RNA for real-time PCR. For fungal biomass analysis, total DNA was isolated from the PM infected leaves. The relative expression of *G. cichoracearum GDSL-*like lipase gene in *Gc* UCSC1 was normalized by the Arabidopsis *GAPCP-2* (*At1g16300*) to indicate the fungal biomass of PM on the infection leaves (Weßling and Panstruga, 2012). The real-time PCR assay was performed using the real-time PCR system. The primers used for RT-PCR and Real-Time PCR are listed in Supplemental Table 1.

## Supplemental data

**Supplemental Figure S1.** Structure-based sequence alignment of R8IP1s from six representative powdery mildew species.

**Supplemental Figure S2.** RPW8.2 associates with the RING domain of GcR8IP1.

**Supplemental Figure S3.** The temporal expression pattern of *GcR8IP1* during powdery mildew infection.

**Supplemental Figure S4.** GcR8IP1-GFP is localized in the nucleus.

**Supplemental Figure S5.** The XVE expression cassette driving GcR8IP1-HA expression in *Arabidopsis*.

**Supplemental Figure S6.** Ectopic expression of *GcR8IP1* compromises immune response in Wa-1.

**Supplemental Figure S7.** Schematic illustration of *GcR8IP1* cDNA.

**Supplemental Figure S8.** Subcellular localization of RPW8.2-GFP in the presence or absence of GcR8IP1.

**Supplemental Figure S9.** GcR8IP1 impairs RPW8.2-triggered H_2_O_2_ production in *N. benthamiana*.

**Supplemental Figure S10.** Ectopic expression of *GcR8IP1* compromises RPW8-mediated resistance to powdery mildew in Wa-1.

**Supplemental Figure S11.** R*P*W8*.2* is induced by BTH but the RPW8.2-YFP remained undetectable *in planta*.

## Acknowledgements

We thank S. Somerville for the *G. cichoracearum* UCSC1 isolate and F. Katagiri for *pad4-1 sid2-1* mutant. This work was partially supported by the National Natural Science Foundation of China (No. U19A2033 to W-M.W. and 32121003 to X-W.C.), the Sichuan Youth Science and Technology Innovation Research Team Foundation (2022JDTD0023 to J.F.), the Natural Science Foundation of Sichuan Province (2022NSFSC0174 to H.W. and 2022NSFSC1699 to Z-X.Z.), and the National Science Foundation (Grant numbers IOS-1457033 and IOS-1901566 to S.X.).

## Author Contributions

W.-M.W., J.-H.Z., S.X., X.-J.W., and X.-W.C. designed experiments and analyzed data; J.-H.Z., Z.-X.Z., and Y.-Y.H. generated and characterized genetic material; J.-H.Z., Y.L., and Z-X.Z. performed cell death and pathogen growth assays; J.-H.Z., J.F., Y.-Y.H., S.-X.Z., G.-B.L. contributed immunoprecipitation and immunofluorescence; J.-H.Z., X.-M.Y., H.W., M.P., B.H., X.-H.H. and J.-W.Z. designed and performed protein-protein interaction assays in yeast and *N. benthamiana*. J.-H.Z., W.-M.W., S.X., and X.-J.W. wrote the manuscript with contributions from all authors.

**Supplemental Figure S1. Structure-based sequence alignment of R8IP1s from six representative powdery mildew species.**

Sequences were aligned using MEGA7 and secondary structures predicted using ESPript (https://espript.ibcp.fr/ESPript/ESPript/). Note that the RING domain (18-61 aa) is almost identical. Blue frames indicate conserved residues, white letters in red boxes indicate strict identity and red letters in white boxes indicate similarity. The secondary structures of β sheets (arrows) and α-helices (spiral) are indicated. QCL08962.1 *Golovinomyces cichoracearum* UCSC1 (GcR8IP1), RKF73030.1 *Golovinomyces cichoracearum* USMG1, RKF64333.1 *Erysiphe neolycopersici*, POS86419.1 *Erysiphe pulchra*, CAD6504834.1 *Blumeria graminis* f. sp. triticale, SZE99581. 1 *Blumeria hordei*.

**Supplemental Figure S2. RPW8.2 associates with the RING domain of GcR8IP1.** Confocal images of BiFC showing the association of RPW8.2-YC with YN-RING transiently co-expressed in *N. benthamiana*. Size bar, 50 μm.

**Supplemental Figure S3. The temporal expression pattern of *GcR8IP1* during powdery mildew infection.**

(A) Quantitative PCR assay showing the *GcR8IP1* expression pattern in *pad4-1 sid2-1* double mutant from 0 to 14 days post inoculation (dpi) of *G. cichoracearum* UCSC1. The data are shown as mean ± s.d. (n = 3), and different letters indicate significant differences (*P* < 0.05) as determined by the one-way Tukey–Kramer test.

(B) Quantification analysis on biomass of powdery mildew from 0 to 14 dpi of *G. cichoracearum* UCSC1. Relative fungal biomass was calculated by the comparison between *G. cichoracearum GDSL-like lipase* gene and *A. thaliana Glyceraldehyde-3-phosphate dehydrogenase of plastid 2* gene (*GAPCP-2*). The data are shown as mean ± s.d. (n = 3), and different letters indicate significant differences (*P* < 0.05) as determined by the one-way Tukey–Kramer test. Similar results were obtained in at least three independent experiments.

**Supplemental Figure S4. GcR8IP1-GFP is localized in the nucleus**. GcR8IP1-GFP was transiently expressed from the *35S* promoter in *N. benthamiana* via the agroinfiltration method. Images were taken at 2 d post infiltration. Size bars, 10 µm.

**Supplemental Figure S5. The XVE expression cassette driving GcR8IP1-HA expression in *Arabidopsis*.**

(A) Quantitative PCR assay showing the relative transcription abundance of *GcR8IP1-RFP*. Expression of *GcR8IP1-RFP* driven by the *35S* promoter in the indicated transgenic lines in Col-0 (WT) background. Error bars represent the mean ± s.d. (n = 3). ND, Not detected.

(B) Immunoblotting analysis on the accumulation of GcR8IP1-RFP in the indicated lines. Total proteins were extracted from 4-weeks-old mature plants leaves. The accumulation of GcR8IP1-RFP was examined by western blotting using an anti-RFP antibody. Ponceau S was used as the loading control.

(C and D) Immunoblotting analysis on the expression of GcR8IP1-HA from the XVE expression cassette in the Col-0 (C) and the Wa-1 background (D). Total proteins were extracted from 4-weeks-old mature plants leaves after treatment of β-estradiol at the indicated time point. The expression of GcR8IP1-HA was examined by western blotting using an anti-HA antibody. Ponceau S was used as the loading control.

**Supplemental Figure S6. Ectopic expression of *GcR8IP1* compromises immune response in Wa-1.**

(A) Representative leaves showing the H_2_O_2_ production in the indicated transgenic lines of XVE-*GcR8IP1*-HA-OE for β-estradiol inducible expression of *GcR8IP1-HA* in the Wa-1 (WT) background at 24 hpi of *G. cichoracearum* UCSC1. The leaves from the indicated lines were spray-treated with β-estradiol or DMSO (mock) 12 h prior to the inoculation of *G. cichoracearum* UCSC1. H_2_O_2_ production was examined by 3,3ʹ-diaminobenzidine staining. Size bar, 1 cm.

(B) Chemiluminescence assay showing elicitation of the oxidative burst induced by chitin in the indicated lines. Leaf disks were punched and incubated with 100 μg/mL of chitin from the 4-week-old plants at 12 h after treatment with β-estradiol or DMSO (mock). Signals were recorded for 30 min after chitin elicit. Error bars represent the mean ± s.d. (n = 8) and asterisks (**, ****) denote significant differences (*P* < 0.01, *P* < 0.001, respectively) as determined by Student ’s *t*-test.

(C and D) Relative expression of the indicated defense-related genes in the indicated lines measured by qRT-PCR. Leaf samples were collected at the indicated time points of inoculation with *G. cichoracearum* UCSC1 after treatment of β-estradiol or DMSO. The data are shown as mean ± s.d. (n = 3), and different letters indicate significant differences (*P* < 0.05) as determined by the one-way Tukey–Kramer test.

**Supplemental Figure S7. Schematic illustration of *GcR8IP1* cDNA.** The indicated inverted repeat was amplified to make the construct for host-induced gene silencing analysis. The primers R8IP1-RT-F and R8IP1-RT-R were used to examine the expression of *GcR8IP1* in qRT-PCR.

**Supplemental Figure S8. Subcellular localization of RPW8.2-GFP in the presence or absence of GcR8IP1.**

(A and B) Representative confocal images showing the subcellular localization of RPW8.2-GFP when it was co-expressed with GcR8IP1-RFP (A), or with 2RFP-NLS

(B) in *N. benthamiana*. Size bar is 50 μm.

**Supplemental Figure S9. GcR8IP1 impairs RPW8.2-triggered H_2_O_2_ production in *N. benthamiana***. A representative leaf showing H_2_O_2_ production triggered by the co-expression of RPW8.2 with GcR8IP1 or with GFP in *N. benthamiana* determined by 3,3ʹ-diaminobenzidine staining at 60 hours post infiltration.

**Supplemental Figure S10. Ectopic expression of *GcR8IP1* compromises RPW8-mediated resistance to powdery mildew in Wa-1.**

(A) Chlorophyll auto-fluorescence images indicate the severity of powdery mildew in the indicated transgenic lines of XVE-*GcR8IP1*-HA-OE for β-estradiol inducible expression of *GcR8IP1-HA* in the Wa-1 (WT) background at 10 d post inoculation (dpi) of *G. cichoracearum* UCSC1. The leaves from the indicated lines were spray-treated with β-estradiol or DMSO (mock) 12 h prior to the inoculation.

(B) Quantification analysis on the chlorophyll fluorescence parameter Fv/Fm from the indicated lines. The data are shown as mean ± s.d. (n = 3), and different letters indicate significant differences (*P* < 0.05) as determined by the one-way Tukey–Kramer test.

**Supplemental Figure S11. RPW8.2 is induced by BTH but the RPW8.2-YFP remained undetectable *in planta*.**

(A) The mRNA expression level of RPW8.2 was quantified by qRT-PCR at the indicated time points after the indicated treatments. Four-week-old R2Y4 plants transgenic for *R82_pro_:RPW8.2-YFP* were spray-treated with 100 μM of the SA functional analogue BTH or DMSO, respectively. Leave samples were collected at time points as indicated. The data are shown as mean ± s.d. (n = 3), and different letters indicate significant differences (P < 0.05) as determined by the one-way Tukey–Kramer test.

(B) Immunoblot detection of RPW8.2-YFP in R2Y4 after Mock or 100 μM BTH treatment. Leaf discs collected at time point as indicate. The Ponceau S-stained Rubisco large subunit was used as loading control.

